# Strong spatial genetic structure in a Baltic Sea herbivore due to recent range expansion, multiple bottlenecks and low connectivity

**DOI:** 10.1101/595629

**Authors:** Pierre De Wit, Per R. Jonsson, Ricardo T. Pereyra, Marina Panova, Carl André, Kerstin Johannesson

## Abstract

In the Baltic Sea, recent range expansions following the opening of the Danish straits have resulted in a low-diversity ecosystem, both among and within species. However, relatively little is known about population genetic patterns within the basin, except for in a few commercially caught species and some primary producers thought to be ecosystem engineers. Here, we investigate the population genetic structure of the ecologically important crustacean *Idotea balthica* throughout the Baltic Sea using an array of 33,774 genome-wide SNP markers derived from 2b-RAD sequencing. We also generate a biophysical connectivity matrix, with which we compare the genomic data. We find strong population structure on small scales across the Baltic Sea, and that genomic patterns in most cases closely match biophysical connectivity, suggesting that current patterns are important for dispersal of this species. We also find a strong signal of multiple bottlenecks during the initial range expansion, in the form of reduced heterozygosity along the historical expansion front. The lack of gene flow among sampling sites in the Baltic Sea environmental gradient potentiates local adaptation, while at the same time also increasing genetic drift in low-diversity areas.

## Introduction

How species manage to colonize a novel environment remains somewhat of a conundrum in evolutionary biology (Bock *et al*. 2015). When we observe nature, examples are plentiful of organisms which have been able to successfully colonize and adapt to environments which are very different from their native ranges (e.g. Hill *et al*. 2011; Cao *et al*. 2016; Barker *et al*. 2017; Frleta-Valić *et al*. 2018). Yet, theory predicts that populations at the edge of a colonization front can accumulate deleterious mutations, the “expansion load” (Peischl & Excoffier 2015), which might become common due to random events such as “allelic surfing” on the expansion wave (Klopfstein *et al*. 2006). Low population sizes at colonization fronts and multiple sequential founder effects should increase the strength of genetic drift, and prevent populations from adapting to novel environmental conditions (Brandvain & Wright 2016). How drift-selection balance acts out along colonization fronts in nature is thus an area of active research (see e.g. Pierce *et al*. 2017; Schrieber & Lachmuth 2017). One pattern that might arise from this process of multiple founder effects along a recently colonized environmental gradient would be a rapid evolution of population structure (Excoffier *et al*. 2009). This would be visible in loci under divergent environmental selection along the gradient, but also in neutral loci due to rapid drift during bottleneck events and reduced gene flow across locally adapted populations (Ibrahim *et al*. 1996). Once the range expansion is concluded, however, the population structure might be diffused again due to population growth and increased gene flow (Hagen *et al*. 2015) – unless there are mechanisms to prevent this from happening, such as assortative mating combined with local adaptation, or if there are strong dispersal barriers in the newly colonized area (Lee 2011).

The Baltic Sea (Figure 1) has recently been applied as an evolutionary model system to study effects of range expansions and adaptation to sharp environmental gradients (Johannesson & André 2006). The connection to the North Sea opened recently, about 8 000 years ago. Previously, the Baltic Sea was a fresh water lake, but since the connection across the Danish Straits opened, a salinity gradient has formed, with a sharp decline in salinity at the entrance to the western Baltic just south of the Danish Straits, followed by a more gentle decline into the Baltic proper and the Bothnian Sea and the Gulf of Finland, where the environment essentially is a freshwater habitat. Populations of marine organisms have since colonized this gradient environment to different extents. These differences have been hypothesized to be due to multiple reasons, for example differences in adaptive (Weinberger *et al*. 2008; Wrange *et al*. 2014), dispersal (Urho 1999; Sjöqvist *et al*. 2015), or reproductive (Jaspers *et al*. 2011) abilities across species, or combinations thereof. In addition to the evolutionary interesting history of the Baltic Sea, it has recently been proposed as a model system for studying future impacts of habitat loss, eutrophication, pollution and over-fishing (Reusch *et al*. 2018), as it is one of the most environmentally impacted seas in the world. Moreover, the Baltic Sea is one of the fastest-warming regions resulting in increasing surface water temperature and reduction of salinity (Meier *et al*. 2012), which is expected to lead to many range shifts with both loss of present biodiversity and introduction of new species (Ojaveer & Kotta 2015). Low functional biodiversity, including genetic diversity, may expose the Baltic Sea to fast ecosystem transformations with potential loss of essential ecosystem services due to low resilience capacity (Österblom *et al*. 2007). In order to predict future biodiversity in the face of ongoing global change, a deeper understanding of population dynamics during range shifts is required (Davis & Shaw 2001).

**Figure 1.**
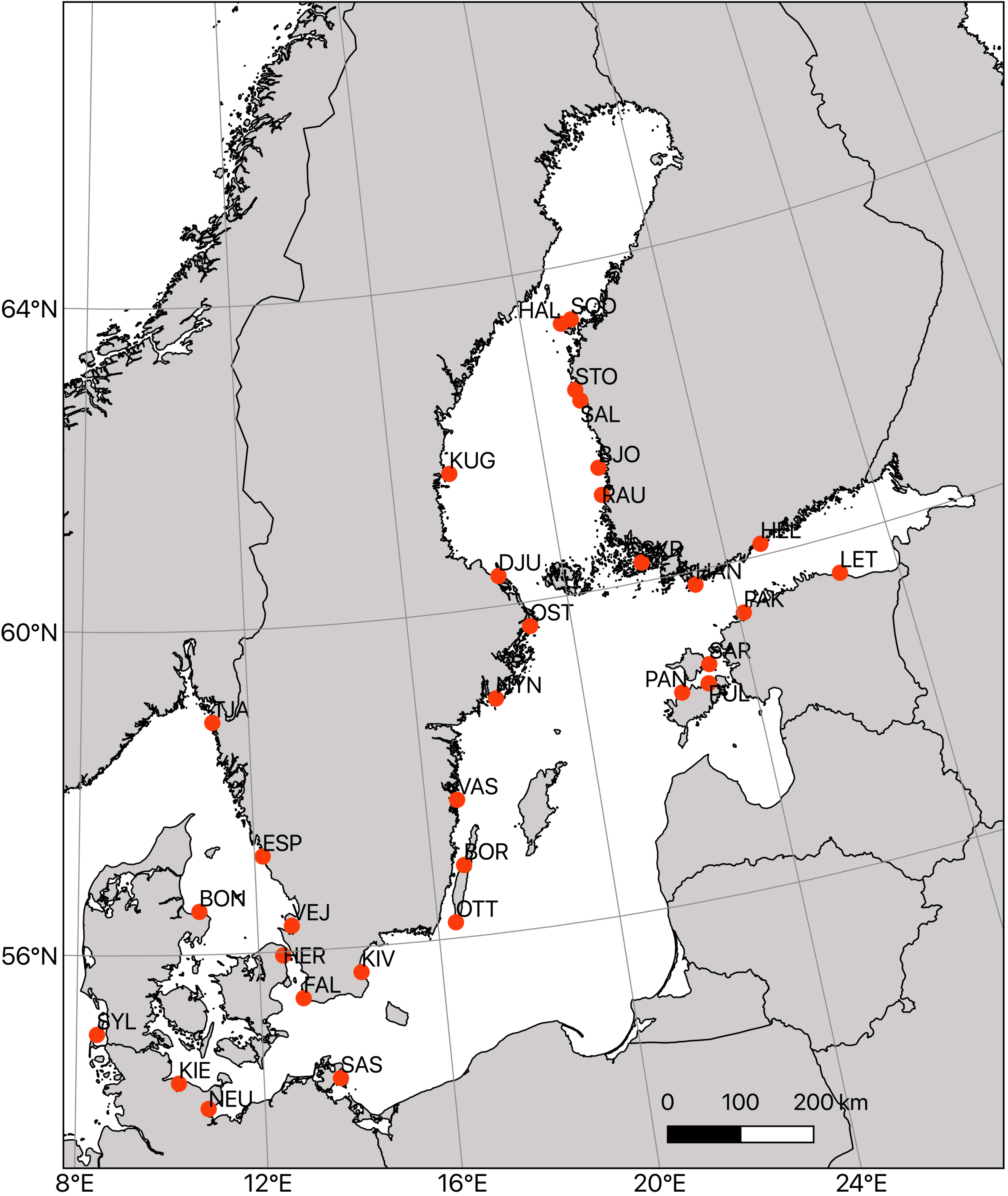
Map of the study area (the Baltic Sea), with collecting sites marked in red.

The Baltic Sea is a multi-basin marginal sea, with rather strong oceanographic barriers separating the different basins (Leppäranta & Myrberg 2009), and there is also limited oceanographic connectivity between Scandinavia and the mainland European coast (Moksnes *et al*. 2014) potentially preventing dispersal of coastal organisms. Thus, we might expect to see signs of reduced gene flow among populations in the Baltic. However, studies of gene flow and population structure are scarce in the Baltic Sea. The studies made to date focus mainly on a few commercially caught fish species (Wennerström *et al*. 2017), where patterns differ substantially. Some fish, such as perch and herring seem to be structured geographically (Olsson *et al*. 2011; Teacher *et al*. 2013), as are species with distinct spawning areas such as cod and salmon (Berg *et al*. 2015; Poćwierz-Kotus *et al*. 2015). However, in other fish, such as three-spined sticklebacks and turbot this is not the case (except in adaptive loci) (Nielsen *et al*. 2004; Guo *et al*. 2015), indicating that population genetic patterns are strongly determined by life history. In addition, a few studies have been made on ecosystem engineering species, mainly the brown algae *Fucus vesiculosus* and *Fucus radicans* (Pereyra *et al*. 2013; Ardehed *et al*. 2016) where population structure has been observed. However, in Baltic Sea *Fucus* species, high incidence of asexual reproduction in the Bothnian Sea confuses the overall pattern of genetic structure (Johannesson *et al*. 2011). These few studies highlight examples of interesting genetic patterns along a colonization front across an environmental gradient such as the Baltic Sea, yet says little about the generality of these patterns. In order to get a better understanding of population genetic mechanisms involved in colonization and range shifts, it is necessary to also study other types of organisms with different life history traits.

*Idotea* is a genus of isopod crustaceans common in the coastal marine environment. Three *Idotea* species have been able to colonize the low salinity conditions of the Baltic Sea in the 8 000 years since the current connection between the North Sea and the Baltic opened (*I. balthica*, *I. granulosa* and *I. chelipes*)(Leidenberger *et al*. 2012). *Idotea* are generalist grazers on different algae and sea grasses, but also have the uncommon ability of being able to survive and grow on a diet of brown algae of the genus *Fucus* (Bell & Sotka 2012). This makes these isopods particularly ecologically important in the Baltic Sea, where the depauperate ecosystem is based on *Fucus* algae as habitat engineers in large parts of the coastal zone, and where *Idotea* is the only grazer on *Fucus*. Indeed, in some areas in the Baltic, densities of *I. balthica* can rise to astonishing numbers and in some cases this can cause a total loss of *Fucus* in an area to grazing (Gunnarsson & Berglund 2012). The three isopod species have colonized the Baltic to different extents, with *I. balthica* going furthest into low-salinity areas in the Bothnian Sea and the Gulf of Finland, followed by *I. chelipes* and *I.granulosa*, respectively (Leidenberger *et al*. 2012), which suggests differences in adaptive capability to low salinity conditions. A recent phylogenetic study of the *Idotea* species in the Baltic suggested that all three species have independently colonized the brackish-water habitat (Panova *et al*. 2016), which might suggest an adaptive pre-disposition. However, the observed differences in colonization might also be due to limitations to gene flow across oceanographic barriers rather than environmental selection. *Idotea* species brood their young, and thus have no larval dispersal phase, which likely make dispersal across the Baltic Sea rare. On the other hand, they are strong swimmers, and they have the potential to raft long distances with free-floating algae (Thiel & Gutow 2005), which could break down connectivity barriers. Investigations of within-species (neutral) population genomic patterns of the isopods in the Baltic Sea might shed light on the respective roles of connectivity barriers and colonization history in shaping the observed distribution of the species, yet no such studies have been done to date.

Here, we perform the first population genomic study on *I. balthica*, and the first study on population genomic patterns across the salinity gradient in the Baltic Sea for a crustacean. We collected isopods from 32 locations spanning from the North Sea to the innermost areas of their distribution range in the Bothnian Sea and the Gulf of Finland. Using 2b-RAD sequencing (Wang *et al*. 2012) we identify ca. 35 000 SNP markers randomly distributed across their genome, which we use to estimate population differentiation, barriers to dispersal, and areas of high- and low diversity. We also estimate a connectivity matrix using a biophysical model based on the oceanographic circulation, which we then compare to the genomic data. We hypothesize that the life history traits, together with the recent colonization of the Baltic Sea have resulted in strong population fragmentation along the Baltic Sea coast. We predict that the population genetic patterns will follow an isolation-by-distance pattern, where oceanographic connectivity will largely explain the genetic patterns. Further, we also predict to see a signal of sequential bottlenecks during the recent range expansion in the population genomic data, in the form of reduced genetic variability further into the Baltic Sea.

## Materials and Methods

Individuals of the three *Idotea* spp. present in the Baltic Sea were collected from 31 locations in the Baltic Sea (including Kattegat/Skagerrak), and in one reference population in the North Sea (Sylt), by snorkelling and picking isopods from *Fucus* plants during August-September of 2014. *Idotea granulosa* and *I. chelipes* individuals were discarded, keeping only individuals of *I. balthica*. Animals were decapitated and stored in 95 % ethanol at −20°C. From most sites, DNA from 20 individuals was extracted, 4 of which twice for technical replication, using a Qiagen Blood & Tissue kit and following the standard protocol, with the addition of RNAse during the last 30 minutes of the tissue lysis step in order to avoid RNA contamination. DNA quantity was measured using a QuBit dsDNA BR assay, and quality was assessed through gel electrophoresis.

### Genotyping

2b-RAD libraries (Wang *et al*. 2012) were prepared from the DNA using a modified version of the laboratory protocol developed by Mikhail Matz, available at: https://github.com/DeWitP/BONUS_BAMBI_IDOTEA. In brief, 100-200 ng of DNA template was fragmented using the type 2b endonuclease enzyme BcgI, after which adapters were ligated to the ends of the excised 36-bp fragments. Fragments were then amplified with barcoded adapters, after which they were pooled equimolarly into 24-sample population pools (20 individuals + 4 technical replicates). All pools were sequenced in an Illumina HiSeq 2500 machine, 50bp single-end, at the Swedish National Genomics Infrastructure’s SNP & SEQ platform at Uppsala University.

All bioinformatic analyses were run on the University of Gothenburg computer cluster ‘Albiorix’ (http://albiorix.bioenv.gu.se/). All commands used in the analyses can be found here: https://github.com/DeWitP/BONUS_BAMBI_IDOTEA. An unpublished draft genome assembly for *I. balthica* was used as a reference for mapping the 2b-RAD data, using bowtie2 (Langmead & Salzberg 2012)(Information for the genome project can be found at https://github.com/The-Bioinformatics-Group/Idotea_genome_project). Then, the standard GATK pipeline (McKenna *et al*. 2010) was followed for SNP calling (using the UnifiedGenotyper), including InDel realignment, quality score recalibration (BQSR) and variant score quality recalibration (VQSR), using sites identically genotyped across all technical replicates as a “True” training set for the machine learning algorithm. Poorly genotyped individuals and sites genotyped at < 80 % of all individuals were filtered out, as well as highly (>75 %) heterozygous sites. The dataset was pruned in order to keep only one SNP site/locus. Finally, technical replicates were discarded from further analysis.

### Summary statistics

Mean pairwise F_ST_, as well as population differentiation p-values using Fisher’s exact probability test, were calculated for all population pairs using GENEPOP (Raymond & Rousset 1995; Rousset 2008). Pairwise F_ST_ were plotted on metric MDS-plots using R. Standard diversity indices (F_IS_, heterozygosity, θ) were calculated using Arlequin v.3.5.2.2 (Excoffier & Lischer 2010). Mean heterozygosity for each population as a response variable depending on environmental factors (summer salinity, winter salinity, summer temperature, winter temperature, MPA status as well as distance from the North Sea as the closest straight line following a coastline) were modelled using a generalized linear model in R. Environmental data for the top 6 m were extracted from the Rossby Centre Oceanographic circulation model (Meier *et al*. 2003) and averaged across 1995-2004 for all stations except for Sylt and Tjärnö, which were outside the domain of the model. For the stations Sylt and Tjärnö empirical data from the ICES database (ices.dk) for depths ≤5 m were averaged for the years 1995-2004 for the nearest sample stations. Summer and winter values were averages for June-August and December to February, respectively. Summer and winter salinity were found to be strongly correlated, and thus winter salinity was removed as a factor.

Also, the asymmetrical distance values “d” (Jost’s D), “gst” (Nei’s G_ST_) and “N_m_” (Alcala *et al*. 2014) were calculated among all population pairs using the divMigrate package in R (Sundqvist *et al*. 2016), within which relative migration networks also were generated, in order to examine the main directions of gene flow in well-connected areas (the Swedish Baltic Sea/Finland coast and the Skagerrak/Kattegat/Danish strait area).

### Principal Components Analysis

An identity-by-state (IBS) distance (1 – IBS) matrix was calculated from the full 33,774 SNP dataset, using plink 1.9 (Purcell *et al*. 2007). A hierarchical cluster dendrogram was produced using the hclust function in R, after which a Canonical Analysis of Principal coordinates (CAP) was performed using the Vegan R package. This method is similar to a PCA, but it rotates the eigenvector axes in such a way as to maximize the among-site differences. Also, a test of the significance of collecting site as a factor was tested through an ANCOVA within the Adonis R package.

In addition, a global F_ST_ outlier analysis was performed using Bayescan v 2.1 (Foll & Gaggiotti 2008). IBS-matrices were calculated separately for outliers and non-outliers, and CAP analyses were performed on the two datasets, in order to determine if outliers had disproportionate or distortive effects on the genetic structure analyses. As outliers and non-outliers showed similar patterns, outliers were not discarded in downstream analyses.

### Geographically explicit population inference

Genotype likelihoods for all SNP sites were extracted from the vcf files using bcftools, which were input to NGSadmix (Skotte *et al*. 2013) for population cluster analysis. NGSadmix uses a probabilistic framework to infer ancestry. K of 2 – 32 was used, with 10 replicates of each. The output q-matrices of NGSadmix were combined for each K and plotted as barplots using CLUMPAK online (http://clumpak.tau.ac.il/index.html). Cross-validation scores were examined for each K to infer the most probable number of population clusters. In addition, separate NGSadmix analyses were run for the Skagerrak/Kattegat/Danish Straits area (K=2-8) and the North part of the Swedish Baltic Sea coast and Finland (K=2-12), as sampling sites in these two areas were non-significantly separated in the CAP analysis described above.

In order to extrapolate ancestry coefficients on geographic scales, the R package TESS (Caye *et al*. 2016) was used. We chose K = 12 for plotting, guided by cross-validation scores. A map of the Baltic Sea was downloaded from https://maps.ngdc.noaa.gov/viewers/wcs-client/, from which a grid was generated with a constraint on elevation from 1 m to −50 m. Finally, ancestry coefficients for the 12 different clusters were geographically imputed and plotted on the map in different colors.

In addition, variability in genetic diversity and effective dispersal were modelled and extrapolated geographically using the software package EEMS (Petkova *et al*. 2015). This software uses an isolation-by-distance model with stepping stones (“demes”) as a null model, and outputs deviations from the model, which can be inferred to be barriers to (or corridors of) dispersal. A full-rank distance matrix was generated by imputing missing genotype values with observed mean genotype at each SNP site, as implemented by the “bed2diffs_v2” tool distributed with the EEMS software. Geographic coordinates from 86 vertices specifying a polygon enclosing the Baltic Sea were extracted from Google Earth (REF), after which the EEMS software was run in three separate mcmc chains, each with default lengths (2 M iterations, burn-in 1 M iterations, writing every 10 K iterations to file), using 300 demes. The output was examined for convergence, and plotted on a map of Europe using the rEEMSplots R package (distributed with EEMS).

### Biophysical model

A biophysical model was used to estimate dispersal probabilities and multigenerational connectivity calculated from stepping-stone dispersal across generations. The biophysical model is based on the oceanographic circulation model NEMO-Nordic that produces water velocity fields with spatial resolution of 3.7 km in the horizontal and 3-12 m in the vertical, and a temporal resolution of 3 h (for details see Hordoir *et al*. 2013). The velocity fields are used by the Lagrangian particle tracking model TRACMASS (de Vries & Döös 2001) to estimate dispersal from a site *i* to a site *j* where results are conveniently summarised as a normalised connectivity matrix with elements specifying the dispersal probability between all sites in the domain (Jonsson *et al*. 2016). The particle tracking model calculated the dispersal from and to 34036 sites where particles were parameterized to mimic dispersal of *I. balthica* assuming the following traits and conditions: reproduction occurs between April to September, drift or swimming of adults or juveniles is in the surface water (0-2 m), the PLD was assumed to be 5 days. We also only considered model grid cells with a mean depth less than 30 m. The dispersal of *Idotea* spp. is poorly known (Leidenberger *et al*. 2012) and the trait combination assumed here possibly leads to an overestimation of realised dispersal. In total, the dispersal simulations of *I. balthica* included 34 million particles.

To identify potential dispersal barriers, we applied a recent clustering method (Nilsson Jacobi *et al*. 2012) based on the connectivity matrix. Identification of subpopulations separated by partial dispersal barriers is here formulated as a minimization problem with a tuneable penalty term for merging clusters that makes it possible to generate population subdivisions with varying degrees of dispersal restrictions. Areas that have an internal connectivity above the dispersal restriction are color-coded, and the transitions of colors thus indicate partial dispersal barriers.

Multigenerational connectivity was calculated from the dispersal connectivity matrix by multiplying the matrix with itself for each generation (Jahnke *et al*. 2018). This procedure calculates the stepping stone dispersal when dispersal probability for all possible routes is summed across all generations for 64 generations. With a mean dispersal distance of approximately 25 km per generation 64 generations should ensure potential connectivity on the scale of the Baltic Sea.

### Genetic vs biophysical connectivity

The correlation between genetic differentiation and multigenerational connectivity estimated from the biophysical model was tested with Mantel tests (Mantel 1967). We used four metrics of genetic differentiation: pairwise F_ST_, and calculated with divMigrate the relative migration using G_ST_, relative migration using D and relative migration using nm. Correlations with both untransformed and logarithmically transformed (log_10_(x+1e-50)) connectivity matrices (64 generations) were tested. Because pairwise F_ST_ data are symmetric we extracted symmetric connectivity matrices by using either the minimum connectivity (minimum of *i* to *j* and *j* to *i*), the maximum connectivity or the mean connectivity. For the asymmetric genetic metrics (from divMigrate) we used the full connectivity matrices. Since most scripts of the Mantel test only accepts symmetric matrices, a script for tests of asymmetric matrices was written using Matlab (v 2018a, Mathworks Inc).

## Results

### Genotyping

The 2b-RAD libraries generated μ=8.64 M reads ± sd 3.72 Mreads, and mapping rates (not including multiple hits) were μ = 20.23 % ± sd 2.26 % (Supplementary data). Using 25,329 loci which had identical genotypes across all replicate pairs as a “true” training set for the VQSR, we estimated the true transition/transversion ratio (T_i_/T_v_) to be 1.348. By examining different truth-sensitivity tranches, we could observe a sharp drop in T_i_/T_v_ at above 95 % truth sensitivity, so we used 95 % as the cut-off tranche for SNP filtering. After filtering for highly heterozygous (likely lumped paralogous) loci, and thinning the dataset to one SNP/ RAD locus, we were left with 57,641 SNPs, and after filtering out poorly genotyped sites, the final genotype dataset consisted of 33,774 SNP sites, genotyped at a minimum of 80 % of 595 individuals (Supplementary data), sampled at the 32 different locations (Figure 1).

### Summary statistics and population differentiation

Pairwise F_ST_ values are summarized in Table 1, above the diagonal, with p-values from Fisher’s exact tests of population differentiation below the diagonal. Due to the high number of loci, p-values are either 1 or approaching 0. Most populations are significantly differentiated from each other, with some exceptions: Within the Skagerrak/Kattegat/Danish straits area, there seems to be ongoing gene flow connecting populations, although the pattern is complex. Along the Swedish south-east coast, neighbouring populations seem connected, with an isolation-by-distance pattern which also connects western Finland to the Swedish populations. Within the Finnish Bothnian Sea coast, the population seems to be panmictic. Finally, the Estonian sampling site at Sarve (SAR) seems to be connected both to Finnish and to Kattegat/Skagerrak populations, although this might be an artefact of the low sample size (10 individuals) found at this site.

**Table 1.**
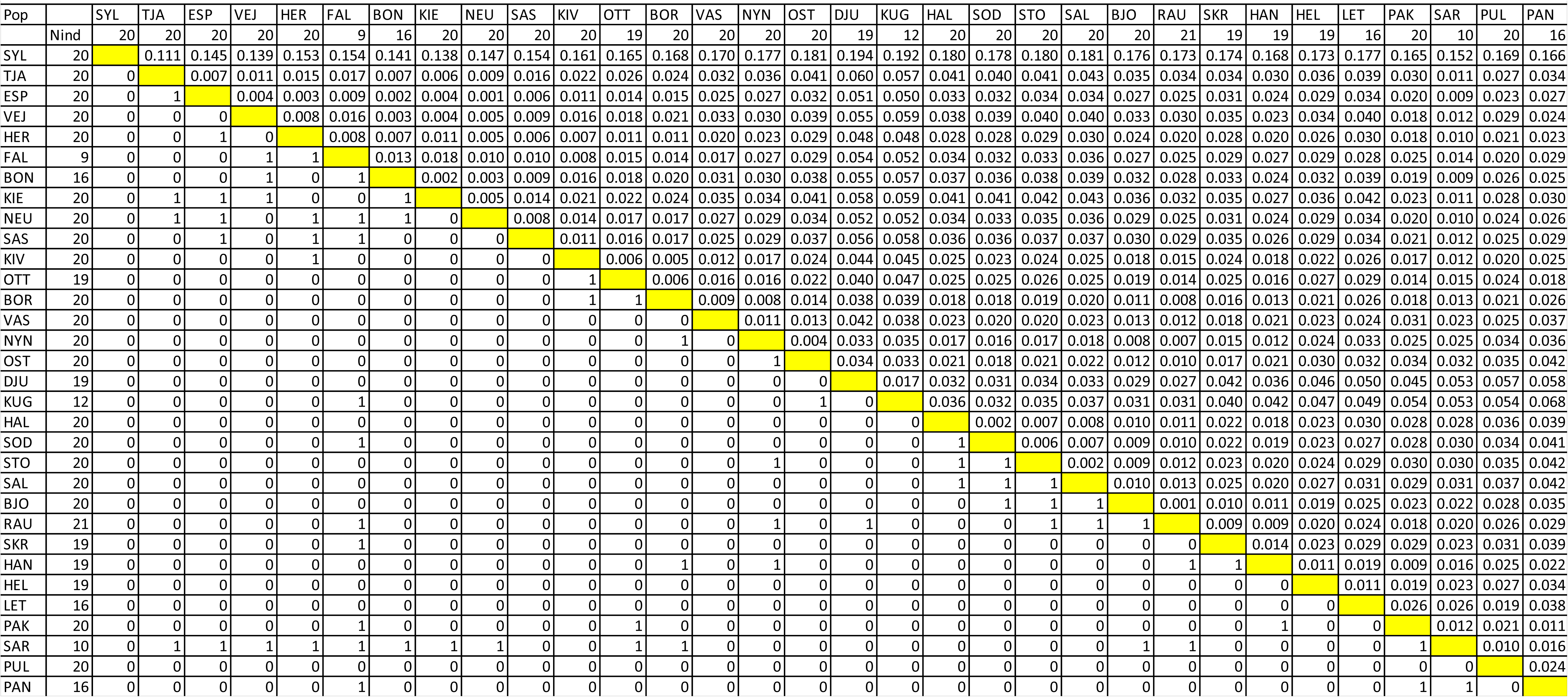
Number of individuals used from each site, pairwise Fst values (above diagonal) and population differentiation Fisher’s exact test p-values (below diagonal; “0” means chi-square value is “Infinity”, and p-value “Highly sign.”).

The pairwise F_ST_ between the North Sea Sylt (SYL) population and all other populations was an order of magnitude higher than others (μ 0.165 ± sd 0.018). Indeed, when plotted in an MDS plot (Figure 2a), SYL is separated from all others along axis 1 which explains 85 % of the variance. When the SYL population is removed, remaining populations scatter more in the plot (Figure 2b), with axis 1 and 2 explaining 61 % and 24 % of the variance in the data, respectively. When SYL is removed, the Swedish Bothnian Sea populations (DJU and KUG) diverge strongly from all others along both of the two primary axes. The great divergence showed in the SYL population caused us to remove these individuals in all further analyses, leaving 575 individuals to be analysed.

**Figure 2.**
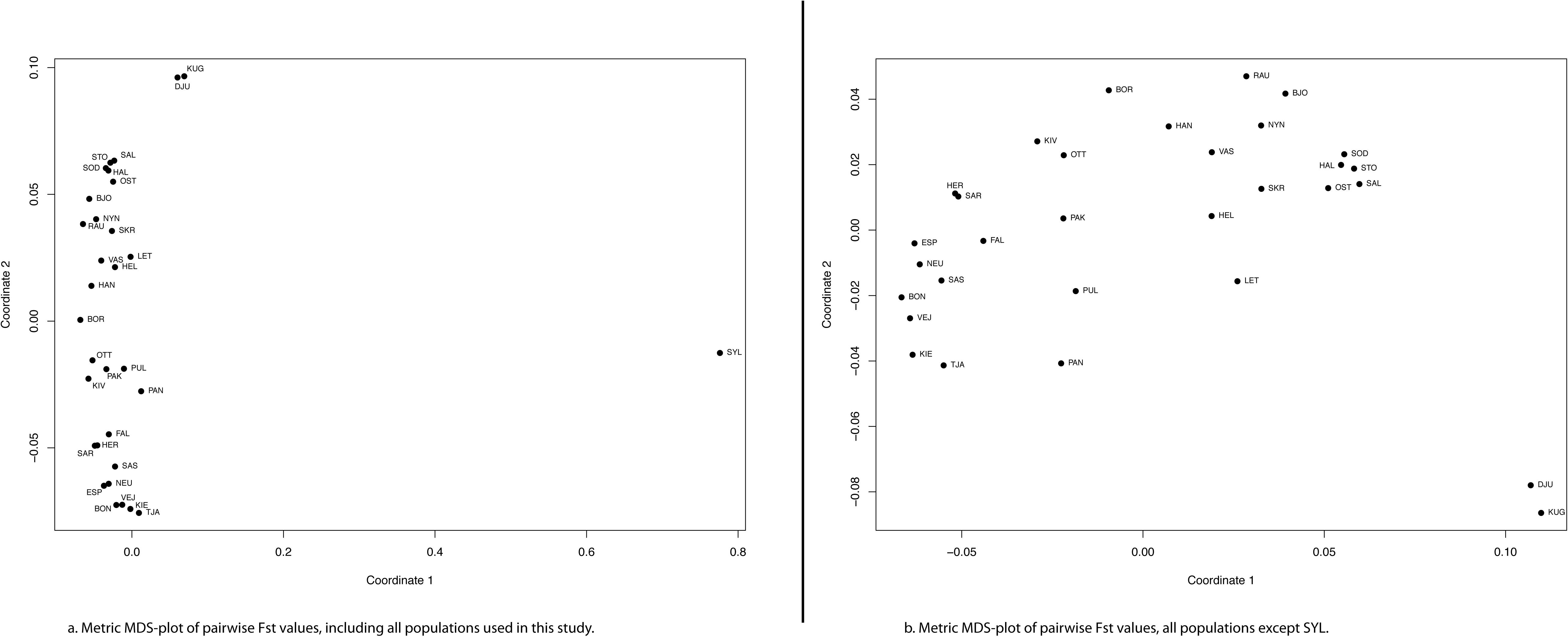
Multi-dimensional scaling plots: a. Metric MDS-plot of pairwise F_ST_ values, including all populations used in this study; b. Metric MDS-plot of pairwise F_ST_ values, all populations except SYL.

F_IS_ values were close to zero in all 32 populations (Supplementary data), and asymmetrical estimates of gene flow (D, G_ST_ and N_m_) indicated that gene flow was roughly symmetrical (±0.05) among most neighbouring sites (Supplementary data). Two notable exceptions to this pattern can be seen, however, along the Swedish Baltic Sea coast, where there is considerably more gene flow in the southerly than northerly direction, and also on the Swedish west coast, where there is more gene flow going north than south.

Mean heterozygosity per collecting site was strongly correlated with salinity (Figure 3, ANCOVA p=8.5e-08), but not with other environmental factors, nor with geographic coastline distance from the North Sea. Heterozygosity was also not determined by if collecting site was a HELCOM Marine Protected Area or not (Figure 3 inset, ANCOVA p = 0.76).

**Figure 3.**
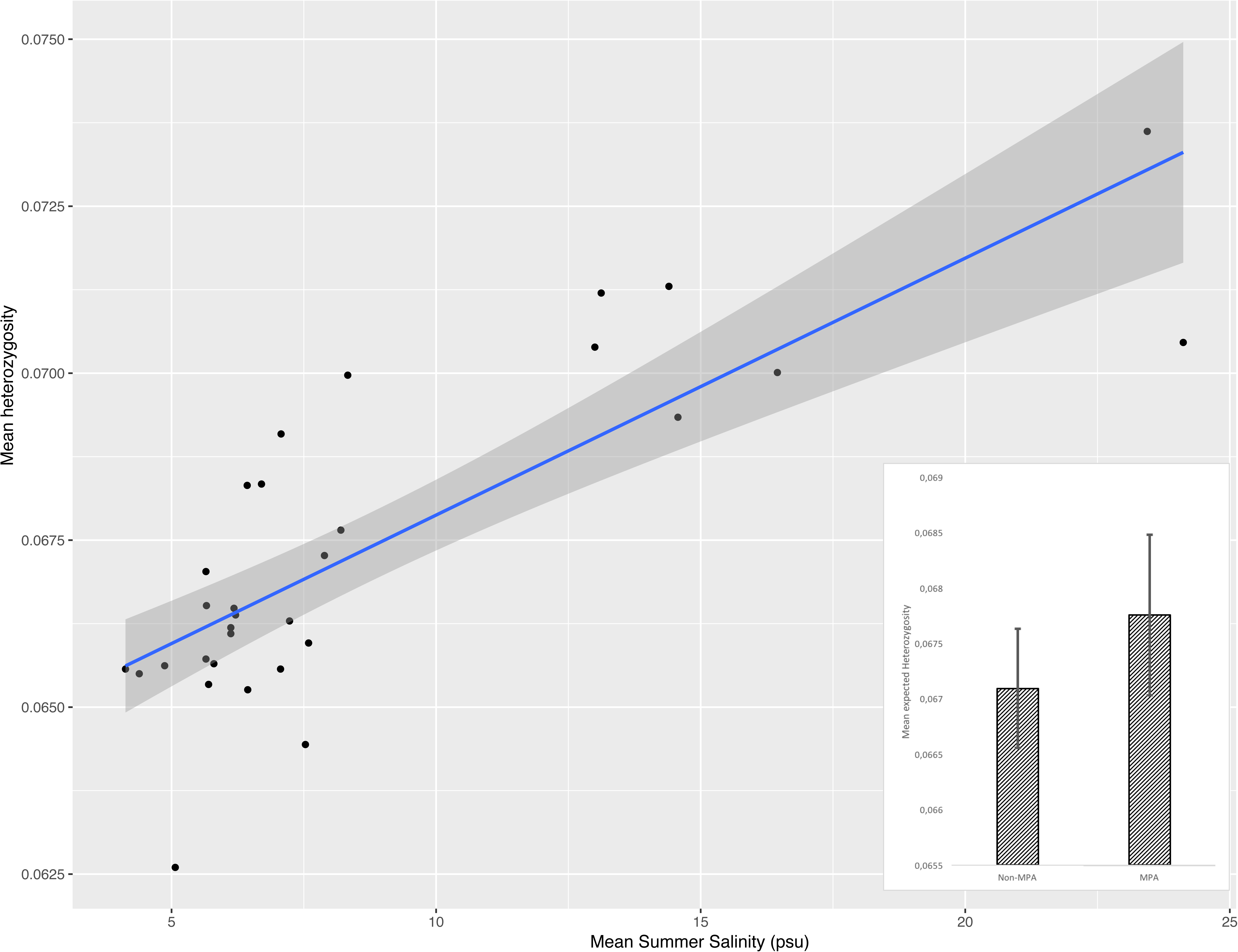
Expected heterozygosity (H_e_) as a function of salinity in the Baltic Sea (p = 8.5e-08). Inset: H_e_ within and outside of MPAs in the Baltic (±SEM; p = 0.76).

### PCA

The hierarchical clustering (Supplementary Fig 1) indicated that some pairs of individuals were closely related to each other. In all cases, the pairs consisted of two individuals from the same site (VAS03/VAS05, PAN18/PAN19, HEL10/HEL16, DJU01/DJU03, DJU18/DJU23, KUG10/KUG13, KUG01/KUG12, HAL02/HAL07, SAL04/SAL09). All of these individuals were well genotyped (Supplementary data), thus it is unlikely that this clustering is an artefact from missing data. Rather, these individuals are likely to be close relatives. In most cases, the pairs of related individuals were sampled in localities with low genetic diversity (see “Summary statistics” section above), indicating that these areas have low effective population sizes.

The CAP analysis (Figure 4) showed strong power to resolve collecting sites from each other, matching geographic patterns almost perfectly, and site was a strongly significant factor, explaining 12 % of the variance (ANCOVA, p=0.001). The outlier analysis identified 487 F_ST_ outliers across all populations (Supplementary Figure 2a), but separate CAP analyses for outliers and non-outliers (Supplementary Figures 2b;c) showed identical patterns, and thus we decided to not exclude outliers. The Skagerrak, Kattegat and Jutland/Western German Baltic Sea coast populations (KIE, TJA, BON, VEJ, ESP, NEU) form one intermixed cluster, while the Öresund (HER) and Eastern German Baltic Sea (SAS) coast are separated from these. Along the Swedish Baltic Sea coast, there is a genetic gradient, with sites being distinct from each other, all the way from FAL (Falsterbo) to OST (Östernäs (Norrköping)). Interestingly, the CAP analysis shows an overlap between the northernmost Swedish Baltic Sea population (OST) and Finnish Bothnian Sea populations, indicating gene flow occurring across the Åland archipelago. In addition, all the northernmost Finnish populations (HAL, STO, SOD, SAL) are intermixed. On the Swedish side, however, the two Bothnian Sea populations (DJU, KUG) seem disconnected from all others. This coincides also with low effective population sizes (see “Summary statistics” section below). The Gulf of Finland populations also form distinct clusters, well separated from each other, both on the Finnish side (HAN and HEL) and on the Estonian side (LET). The Estonian PAK (Pakrinieme) site, located at the entrance of the Gulf of Finland, distinguishes itself from the other Gulf of Finland populations along CAP1, clustering closer to western Estonian individuals (although separated along CAP2), sites are also well distinguished, apart from PAN and PUL overlapping each other, and the SAR population showing a large variance in eigenvalues, with a few individuals scattering closer to the German populations.

**Figure 4.**
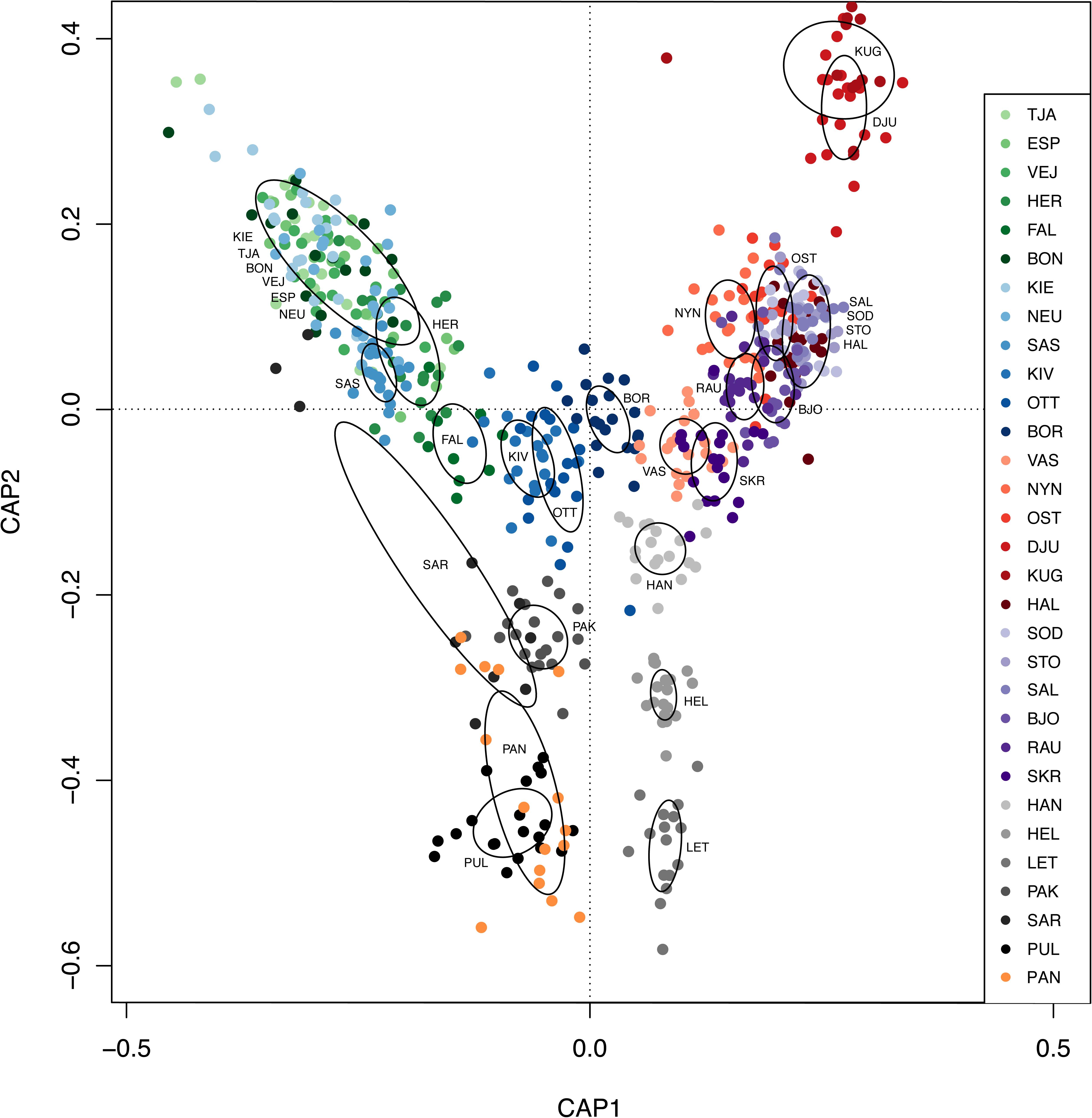
Constrained ordination plot of identity-by-state distances among all *I. balthica* individual used in this study. Circles represent 95 % confidence intervals for collecting sites.

### Spatially explicit population structure

The admixture analysis was run for K = 2 – 32 (Supplementary Figure 3), and examining the cross-validation scores (Supplementary Figure 4), it was clear that a plateau was found at K = 11 – 12, although the software did identify more fine-scale population structure also at higher K values. Thus, we chose to focus on K = 11 and 12 (Figure 5). K = 11 and 12 only differ in that the TJA population (closest to the North Sea) separates out from remaining western Baltic sites, while it clusters together with them at K = 11. The observed pattern is similar to that of the CAP analysis above, where Western Baltic sites cluster together, as do Finnish Bothnian Sea, and Gulf of Finland ones (Including the Estonian LET site), while an isolation-by-distance pattern can be seen along the Swedish south coast. Western Estonian sites clearly separate out from all other sites. Separate admixture runs for the well-mixed areas (Western Baltic, not including FAL, and also the Swedish Baltic Sea/Finland area) found K = 3 to be optimal in the western Baltic and K = 4 to be optimal in the east (Figure 5). In the west, HER and SAS cluster together, TJA is separate from others, while all other sites group together. In the east, there is one large cluster all along the Northern Bothnian Sea coast of Finland, one cluster in the Gulf of Finland, and two on the Swedish coast (NYN, OST) and VAS, respectively, while the geographically intermediately located sites RAU and BJO seem to contain a mix of genetic material from the different groups.

**Figure 5.**
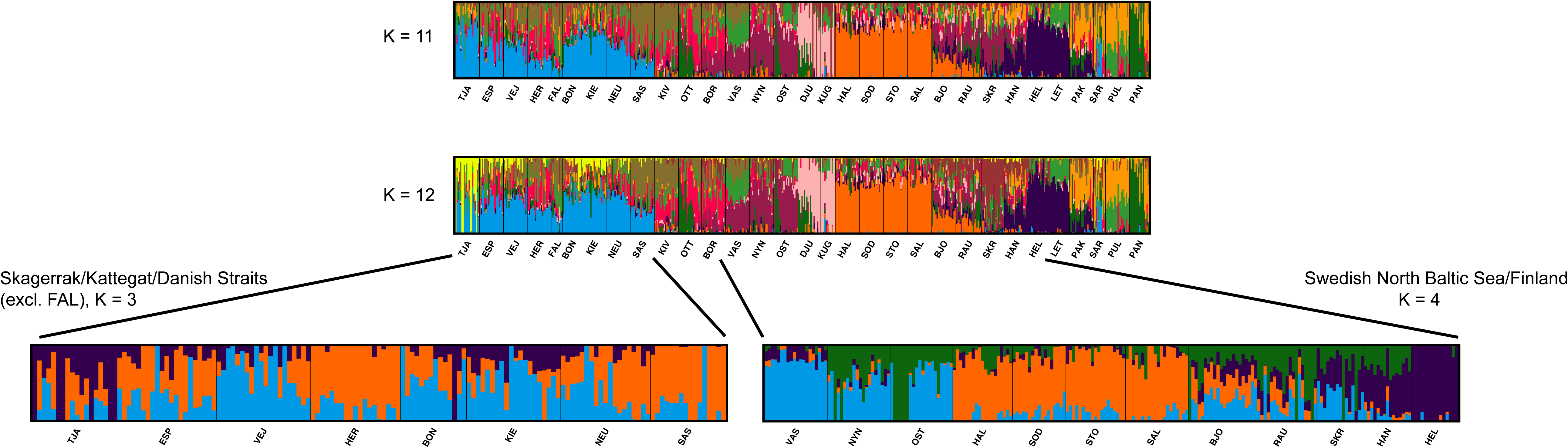
Admixture plots of the entire dataset (two top rows) with K = 11 and 12, and separated into the Skagerrak/Kattegatt/Danish straits areas (K = 3; bottom left) and Finland/Swedish north Baltic (K = 4; bottom right).

The results at K = 12 were imputed on a map of the Baltic Sea, in order to clearly illustrate the geographic distributions of the different genetic clusters, in order to facilitate potential future management decisions (Figure 6). Admixture coefficients are illustrated in different color scales, with q approaching 1 in dark shades, and lower scores in progressively lighter ones. Regions with steep color gradients can be thought of as areas with dispersal barriers, separating genetically divergent populations.

**Figure 6.**
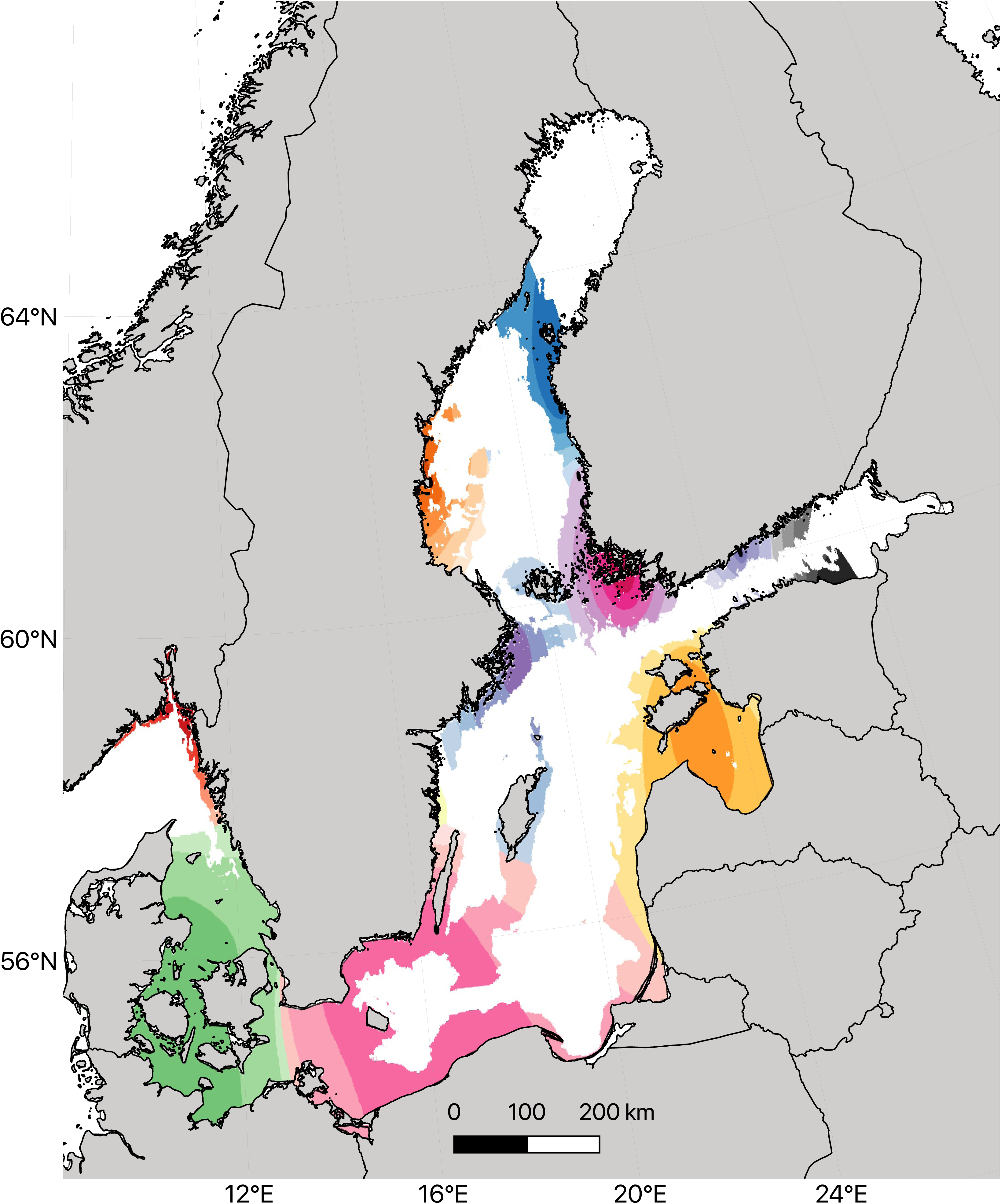
Admixture coefficients interpolated geographically, with population clusters in different color scales (K = 12).

In addition, deviations from stepping-stone isolation-by-distance (IBD) patterns were inferred geographically. Deviations from IBD are indicative of barriers to (and corridors of) dispersal. Genetic diversity (q) was higher in the Skagerrak area and lower in the Baltic Sea, as expected, with particularly low values on the Swedish Bothnian Sea coast and in western Estonia (Figure 7a). Effective dispersal (m) inference (Figure 7b) indicates a corridor of gene flow from northern Denmark to the north part of the Swedish west coast, and one corridor connecting the north and south coast of the Gulf of Finland (HEL and LET). Although these two sites are separated in the CAP plot (Figure 4), a large proportion of their loci do cluster together in the admixture analysis (Figure 5, purple). Surprisingly, a corridor of gene flow was also found to connect southern Sweden with western Estonia, through Poland, Latvia and Lithuania. While this might be an artefact of the non-existent sampling scheme in this part of the Baltic Sea, the genetic similarity between some individuals collected in SAR (Estonia) to southern Sweden seen in the CAP plot (Figure 4) could also be seen as an indication of recent gene flow.

**Figure 7.**
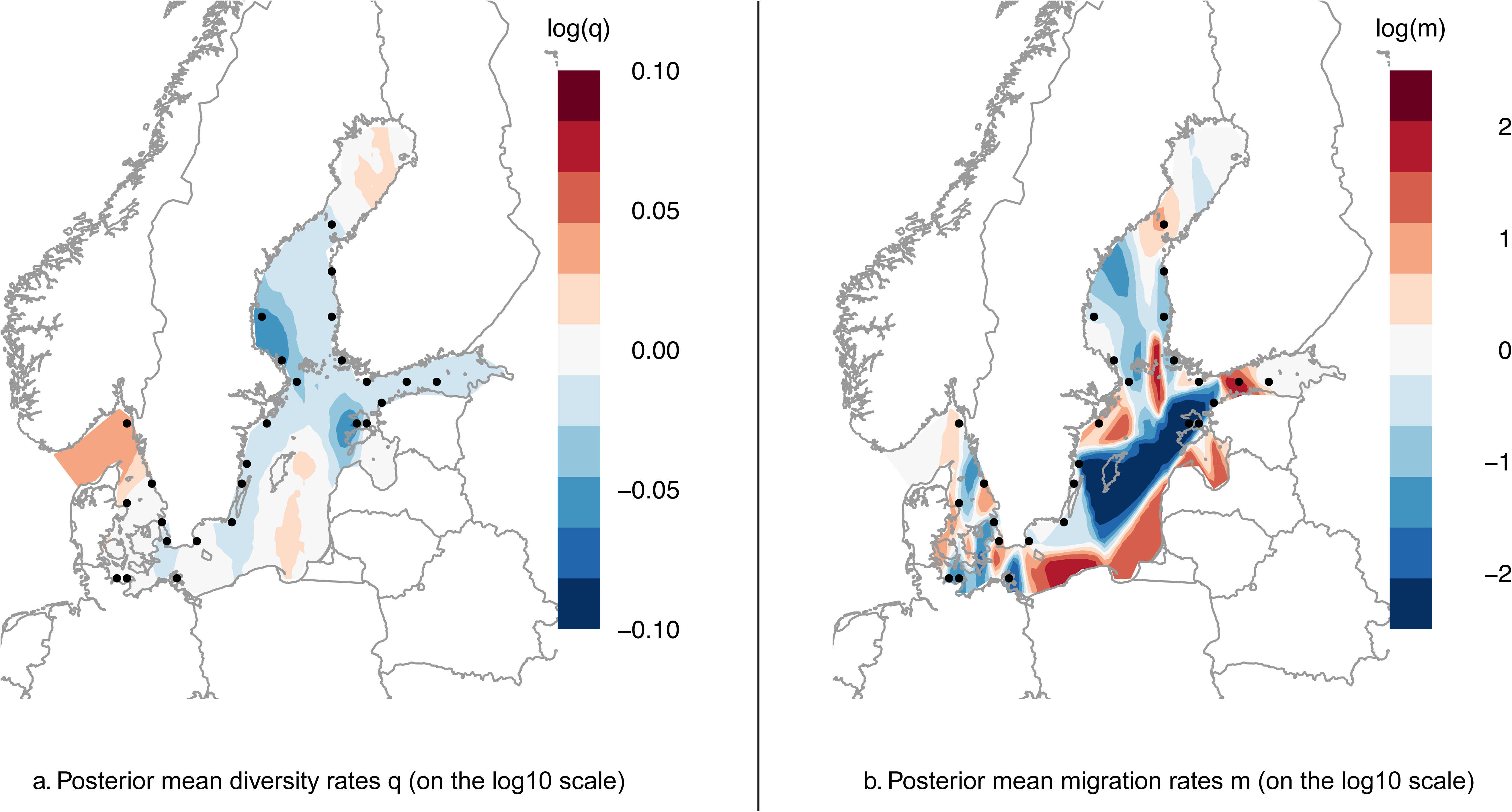
Deviations from isolation-by-distance imputed geographically as: a. Posterior mean diversity rates q (on the log10 scale); b. Posterior mean migration rates m (on the log10 scale).

### Biophysical model

The multigeneration connectivity matrix (64 generations) based on the biophysical particle model (Figure 8; Supplementary data) shows that there are certain regions with high internal connectivity, which are less well-connected to each other. The western Baltic, the Finnish Bothnian Sea, southern Finland and western Estonia are four such regions. Along the Swedish south-east coast, an isolation-by-distance pattern can be seen. Connectivity is in general symmetrical, with a few interesting exceptions: There is potential gene flow from the Finnish coast (SKR/BJO/RAU) to the Swedish East coast (NYN/OST), but not in the other direction. Instead there is potential flow from the more northerly DJU locality in the Swedish Bothnian Sea to the Finnish sites. Along the Finnish Bothnian Sea coast, connectivity also seems to be asymmetrical, with more flow going northward then southward. Interestingly, the particle model also indicates high potential connectivity from the western Baltic to western Estonia, via Poland, Lithuania and Latvia. Also, from the Swedish south coast (FAL, KIV) there is potential connectivity to the western Estonian sites.

**Figure 8.**
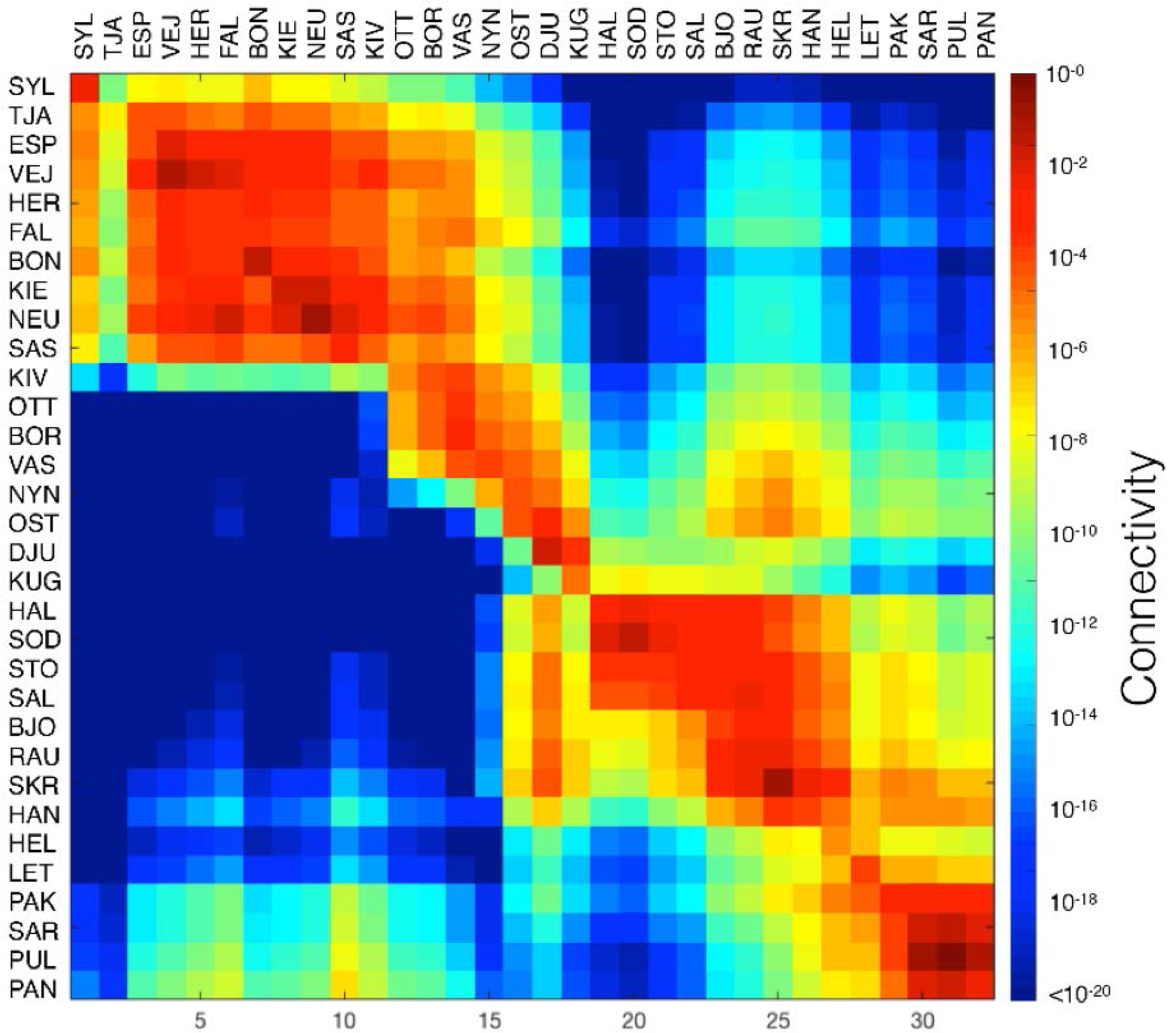
Biophysical connectivity matrix among the sites used in this study, based on a multigenerational iteration of the particle tracking model.

The barrier analysis, which is a way to project the multigeneration connectivity matrix in a geographic dimension, also identified regions with high internal connectivity within the Baltic Sea, which were separated by barriers with high resistance to dispersal (Figure 9). The number of regions varied according to the selected threshold of allowed mean dispersal among regions, and our model with the lowest cut-off value (Figure 9D) most closely matches our genetic data, in that it identifies the separation between the Swedish Bothnian Sea population and others. However, this cut-off value also separates northern from southern Gulf of Finland populations, which we do not see in the genetic data; here the barriers in Figure 9C better represent the genetic data. The barrier analysis averages connectivity values to and from sites, and does thus not well represent instances of highly asymmetric gene flow, which could explain this discrepancy.

**Figure 9.**
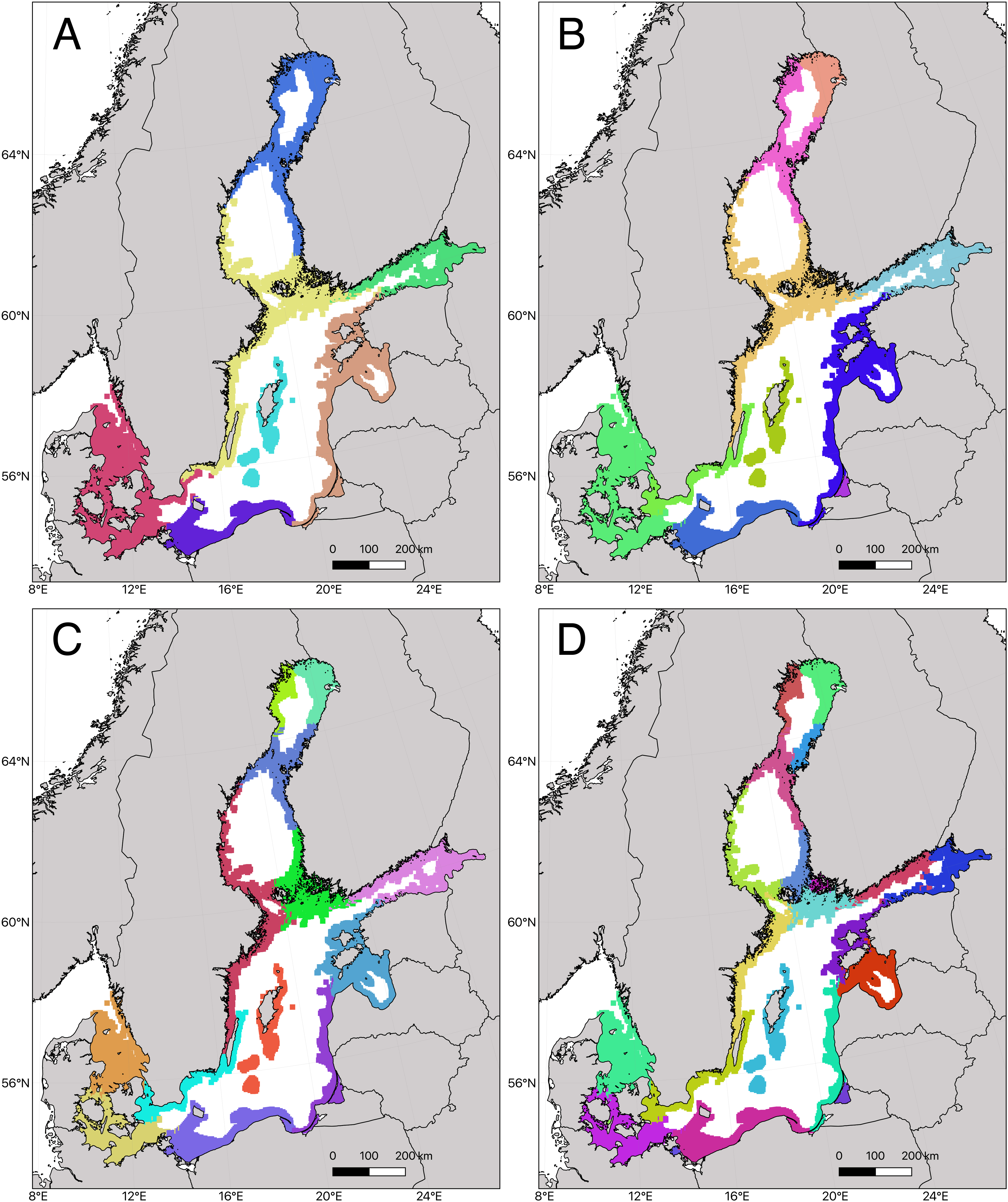
Connectivity barrier inference, clustering areas of high connectivity marked as different colors. Threshold for barrier identification ranges from low (a) to stringent (d).

The biophysical model (mean connectivity) is strongly correlated with pairwise genetic distance (F_ST_) (Mantel p < 1e-5, R= −0.45). Asymmetrical distance values provide a slightly lower fit (N_m_ R = 0.41, G_ST_ R = 0.41, D R = 0.39) but are still strongly significantly correlated to the biophysical connectivity (p < 1e-5 for all three). Interestingly, a simple measure of geographic distance (measured along the shortest coastline connecting two sites) is also significantly correlated to the pairwise F_ST_ matrix (R = 0.41, p< 1e-5).

## Discussion

The population genomic data produced here paints a clear picture of a highly structured meta-population of *I. balthica* in the Baltic Sea. In general, gene flow even between neighboring sites seems to be low, sharply contrasting with patterns in fish species such as sticklebacks (DeFaveri *et al*. 2013). Further, the genetic patterns match the biophysical connectivity model closely, indicating that limitations to dispersal are the most important factor affecting genetic structure in this species. As this species broods its young, there is no pelagic larval dispersal. Adult animals have the potential for long-distance dispersal (mainly through rafting)(Thiel & Gutow 2005; Clarkin *et al*. 2012; Winston 2012), but it seems that *I. balthica* do not do this to any great extent in the Baltic Sea (Leidenberger *et al*. 2012). One potential reason could be the high predation risk – *Idotea* spp. are usually found in high numbers only in thick algal belts, where they are well camouflaged, and they are a favored food source for multiple fish species (Merilaita 2001). So, any long-distance swimming or rafting with small algal fragments would entail a high risk of being eaten. The biophysical model of connectivity included a short pelagic duration of 5 days and a habitat delimited by areas more shallow than 30 m. Although this may overestimate the dispersal capability of *I. balthica*, the model suggests several areas with high resistance to gene flow as visualized in the barrier analysis (Figure 9). The barrier analysis may be viewed as the sequential breaking up into well-connected regions as the threshold of allowed inter-regional dispersal is continuously relaxed (Nilsson Jacobi *et al*. 2012). Some dispersal barriers are consistent across a range of dispersal thresholds (e.g. between western Estonia and the Gulf of Finland), whereas other barriers are more labile (e.g. the Swedish east coast). In general, the barrier analysis well predicted the population structure although additional factors, not considered in the biophysical model, e.g. wave exposure and habitat distribution may further modify the resistance to gene flow. There are a few exceptions to the correlation between the biophysical model and the genetic data. One of the most striking genetic patterns is that the Swedish Bothnian sea populations (DJU and KUG) seems to be strongly isolated from all other sites, while the biophysical model indicates that dispersal can occur along the Swedish coast except in the most stringent setting of the barrier analysis (Figure 9D). This is obvious both from pairwise F_ST_ values and from the low genetic diversity in these areas. However, when also the possibility for asymmetric gene flow is considered (heat-map in Figure 8), the biophysical model shows that DJU and KUG are almost isolated from incoming gene flow, although they are expected to act as sources to other locations (e.g. in eastern Finland). It seems that the Åland archipelago and the current patterns in this area provide an effective barrier to westward gene flow, instead redirecting dispersal from Sweden to the Finnish archipelago sea, and then northward along the Finnish Bothnian Sea coast, rather than along the Swedish Bothnian Sea coast. Another striking feature is that while the biophysical model indicates potential dispersal from Finland to Sweden in the northernmost part of the Bothnian Sea (the “Kvarken” area), and that *I. balthica* is quite common on the Finnish side (SOD/HAL), we do not find any isopods on the Swedish side. In fact, our Kuggören (KUG) sampling site is the most northern point of the species distribution on the Swedish coast. There is plenty of available habitat (*Fucus* belts) also to the north of this site, although not continuous all along the coast, and a recent niche modelling study (Leidenberger *et al*. 2015) also showed that the northern Bothnian Sea coast should be inhabitable to the isopods. However, perhaps the environmental conditions (low salinity/high ice cover), fragmentation of *Fucus* belts, and the lack of available habitat in the Kvarken area restrict dispersal from Finland to Sweden, while the prevailing southerly current along the coast effectively limits the isopods’ ability to disperse northward along the Swedish coast. Although these animals are tolerant to low salinity conditions, there is a limit beyond which the animals cannot survive (Leidenberger *et al*. 2012; Rugiu *et al*. 2018; Kotta *et al*. 2019). Before this point, however, reproduction might be affected due to low fertilization success (Vuorinen *et al*. 2015).

The Finnish Bothnian sea coast, on the other hand, is one large mixed population, with gene flow mainly occurring from the south towards the north. As prevailing currents move northward along the Finnish coast, this is perhaps not so surprising. It might be speculated that the northernmost edge of the Finnish population might have low reproductive output, which would explain the lack of dispersal to the Swedish side, while at the same time maintaining high population density due to constant input from more southerly areas. The strong genetic separation of the Swedish and Finnish Bothnian Sea coasts can be contrasted to genetic patterns observed in *Fucus*, where *F. radicans* is well-mixed throughout the region, whereas *F. vesiculosus* is subdivided both among countries and also among sampling sites within both countries (Pereyra *et al*. 2013). This indicates that different processes are involved in structuring the algal populations from the herbivores; an interesting future avenue for research would be to more closely compare current gene flow and colonization history at the two trophic levels.

The Estonian coast is divided into two distinct regions, one in the Gulf of Finland (with populations connected to Finnish Gulf of Finland populations) and western Estonia with the islands of Hiiumaa and Saaremaa. Interestingly, pairwise F_ST_ (Figure 2), PCA (Figure 4) and the migration surface analysis (Figure 7) all indicate gene flow from southern Sweden and eastern Germany (SAS) to Estonia (SAR site). Although the biophysical model indicates a relatively high potential dispersal in this direction (Figure 8), the Baltic Sea coastline of Poland, Latvia and Lithuania is sandy and lack *Fucus*, which should provide an effective barrier to gene flow. Thus, the observed genetic connection along this coast could be an artefact of the lack of sampling points in this area. Or, it is possible that the isopods can use *Zostera* beds in this area as stepping stones for dispersal. In order to illuminate the patterns of gene flow in this area, further sampling in *Zostera* beds will be necessary.

Along the Swedish coast, starting from the Öresund (HER) site, we see a clear linear isolation-by-distance pattern, with strong population divergence, asymmetric gene flow from the west to the east (D, G_ST_ and N_m_; Supplementary data), and with sequential reductions in genetic diversity (heterozygosity) further in to the Baltic. This is clear evidence of the colonization front moving into the Baltic Sea along the coast, with multiple founder effects. This area would be ideal to investigate further for the demographic history and the timing of the range expansion, as it also crosses the strongest salinity gradient in the Baltic. Here, it might be possible to tease apart the genetic effects of natural selection and demographic effects of a range expansion in the future, in order to gain a deeper understanding of the interplay between selection and drift along a range expansion. The lower heterozygosity in the Baltic proper corresponds to a 12 % drop compared to the most diverse population on the Swedish west coast (TJA). Interestingly, this corresponds very well to the average drop in Baltic nuclear genetic diversity (11-12%) found in the meta-analysis performed by Johannesson & André (2006).

We also examined if there were any clear differences in genetic diversity within and outside of HELCOM marine protected areas, and found no significant differences. This lack of effect of MPAs on genetic diversity has been documented also in other species in the Baltic Sea (Wennerström *et al*. 2017), although there are documented positive effects in other areas of the world (Lester *et al*. 2009). This could be due to the lack of commercial harvest affecting *I. balthica*, the fairly recent implementation of MPAs, and the absence of relevant management plans. According to both the United Nations Convention of Biological Diversity (CBD 1993) and the European Union Marine Strategy Framework Directive (Directive 2008/56/EC), designation of protected areas should be planned with regard to conservation of biodiversity, including within-species genetic diversity (Laikre *et al*. 2016). On the Baltic Sea scale, the Helsinki Commission also recognizes within-species diversity as an important factor in determining ecosystem resilience (HELCOM 2009). However, this has to date not been included in management plans of MPAs in the Baltic Sea (Laikre *et al*. 2016), but hopefully new knowledge about population genetic patterns will help support management efforts in the future. A recent study (Sandström *et al*. 2019) found that a major reason for local managers not using genetic diversity in day to day operations, is that they find this type of information difficult to interpret. We thus here have tried to provide easy-to-interpret maps of genetic diversity (Figure 7) and population subdivision (Figure 6), which we hope will be useful for managers, e.g. in defining management units. One important such would be the Swedish Bothnian Sea coast, which hosts an isolated population of small size. Establishment of MPAs in that area would probably be very beneficial to the coastal animals there. Updated management advice will be continually posted on the BAMBI project website at: https://bambi.gu.se/baltgene.

In this study, we identify 487 global F_ST_ outliers. Population structure in outliers mirror those in putatively neutral loci, indicating that the range expansion and natural selection processes have acted in concert. However, it might be possible to use the independent colonization events into the Bothnian Sea and the Gulf of Finland, in order to identify loci involved in adaptation to low salinity. An interesting avenue for further research would thus be to perform association tests of allele frequencies at individual loci with environmental parameters (primarily salinity), while controlling for the neutral population structure. In addition, further experiments linking genotype and phenotype would be necessary in order to gain more conclusive evidence concerning genes involved in salinity adaptation in the Baltic Sea. Although we currently do not possess knowledge of the function of outlier loci, it is possible that future genomic information, when available, could help with annotation. Identification of alleles involved in local adaptation will be relevant for predicting range shifts in the face of ongoing climate change. A significant warming and decrease in salinity is expected for the Baltic Sea until the end of this century. Considering the apparent poor dispersal ability of *I. balthica* it may be difficult to track the receding salinity gradient by locally adapted genotypes resulting in wide local extinctions. Consequently, it will be important to monitor the northern distribution limit of this important grazer in the future as the environment continues to change, and conservation efforts might need to be focused on facilitating dispersal southward along the salinity gradient, especially along the Swedish Bothnian Sea and northern Baltic Sea coasts.

## Acknowledgements

This work resulted from the BONUS BAMBI project was supported by BONUS (Art 185), funded jointly by the EU and the Swedish research council FORMAS. We would like to thank the Swedish National Genomic Infrastructure (NGI) and the SNP&SEQ Technology platform at Uppsala University for sequencing. All bioinformatics analyses were run on the Albiorix computer cluster (http://albiorix.bioenv.gu.se/) at the Department of Marine Sciences, University of Gothenburg. Finally, we would like to thank our collaborators Melanie Heckwolf, Britta Meyer, Luca Rugiu, Veijo Jormalainen, Eva Rothäusler and Merli Pärnoja for assistance in field campaigns.

## Supplement

**Supplementary data.** Excel document containing mapping statistics, F_IS_ values for each population, asymmetric genetic distance matrices (N_m_; G_ST_; Joost’s D), and the biophysical connectivity matrix from the particle tracking model.

**Supplementary Figure 1.** Hierarchical clustering dendrogram of identity-by-state distances among all individuals used in the study.

**Supplementary Figure 2.** Results from outlier analysis: A. F_ST_ values plotted against q-values with FDR (vertical line) = 10e-4; B. CAP-plot of 487 F_ST_ outliers only; C. CAP-plot of non-outliers only.

**Supplementary Figure 3.** Admixture barplots with K ranging from 2 to 32.

**Supplementary Figure 4.** Cross-validation scores of admixture output, with K ranging from 2 to 32.

