## Supplementary figures and images for "Strong spatial genetic structure in a Baltic Sea herbivore due to recent range expansion, multiple bottlenecks and low connectivity"

### Supplementary Figure 1

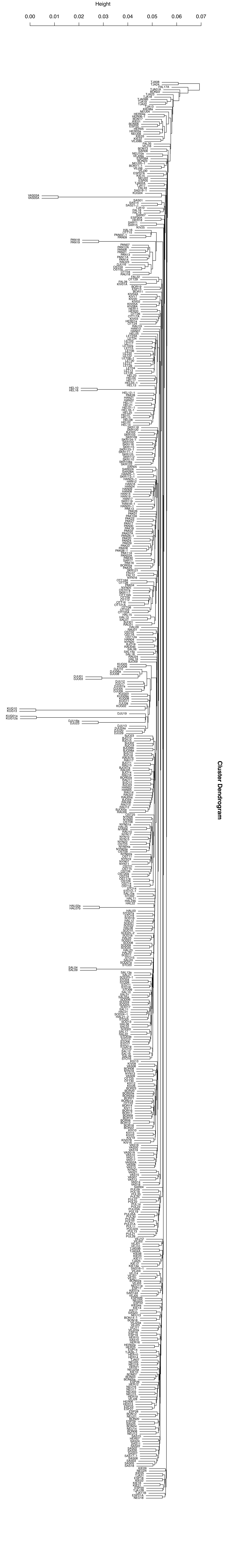

### Supplementary Figure 4

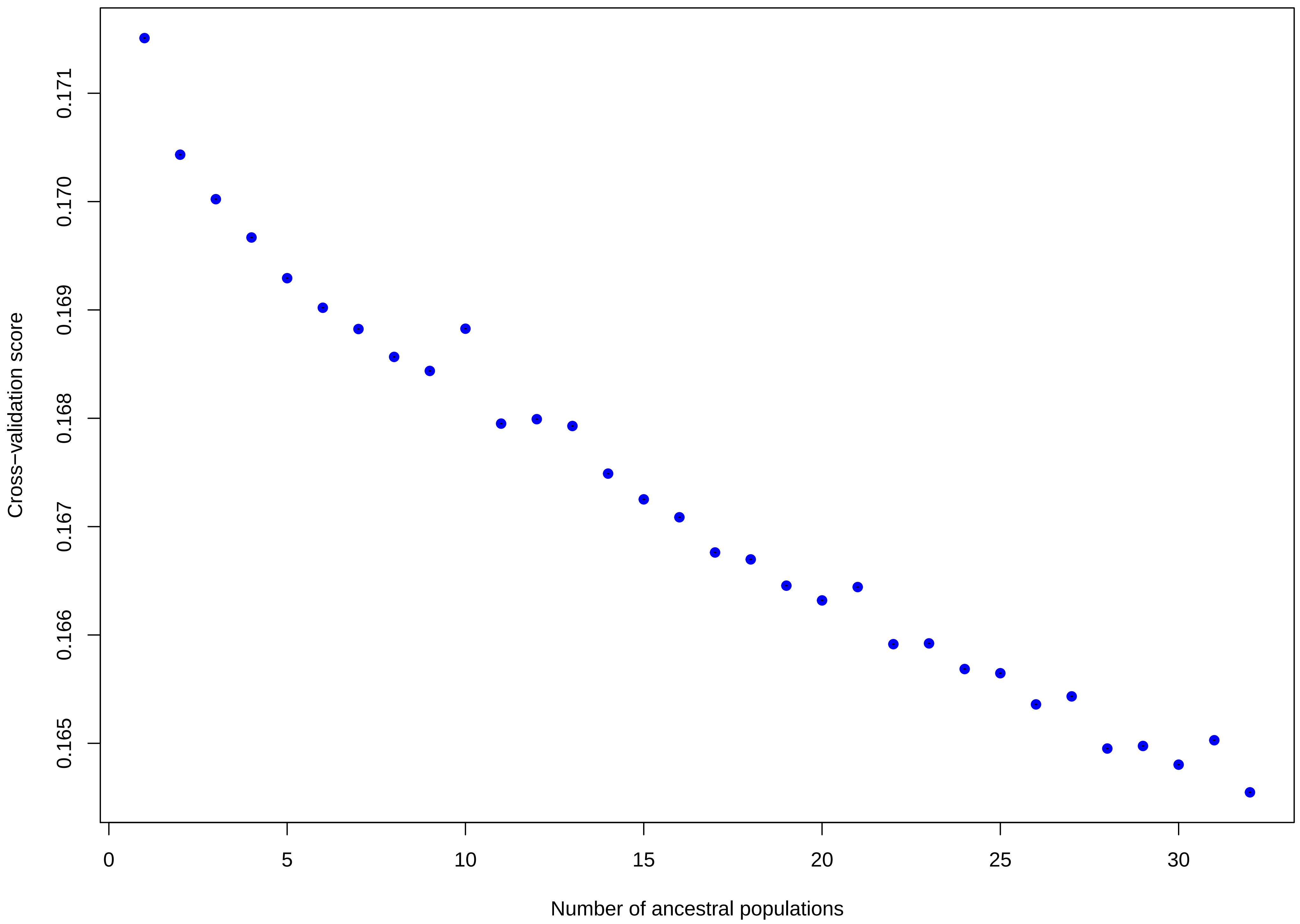
