## Supplementary Figure 2 for "Strong spatial genetic structure in a Baltic Sea herbivore due to recent range expansion, multiple bottlenecks and low connectivity"

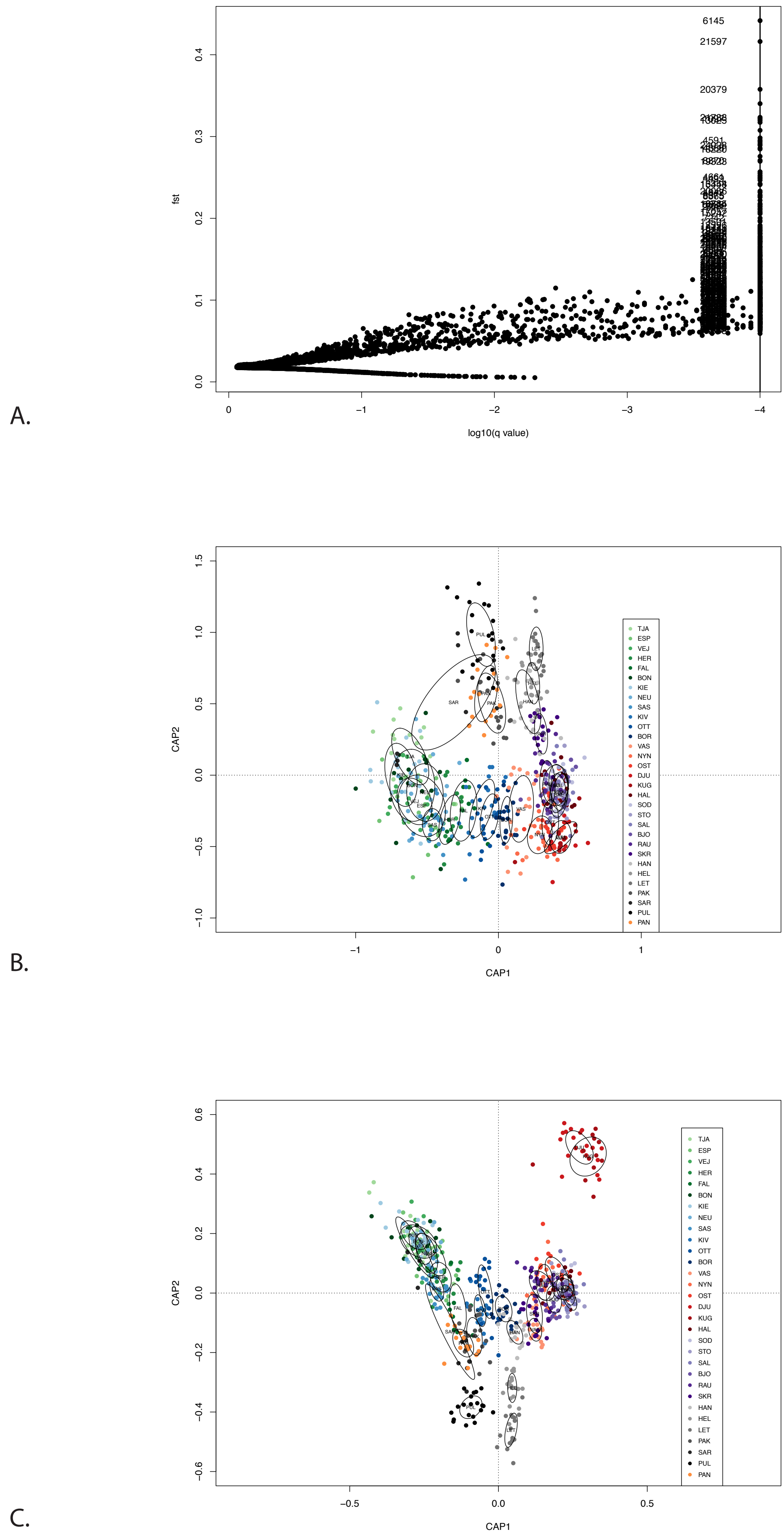


**Supplementary Figure 2**. Results from outlier analysis: A. Fst values plotted against q-values with FDR (vertical line) = 10e-4; B. CAP-plot of 487 Fst outliers only; C. CAP-plot of non-outliers only.
