## Supplementary Figure 3 for "Strong spatial genetic structure in a Baltic Sea herbivore due to recent range expansion, multiple bottlenecks and low connectivity"

### CLUMPAK main pipeline - Job 1548418484 summary

#### Major modes for the uploaded data:

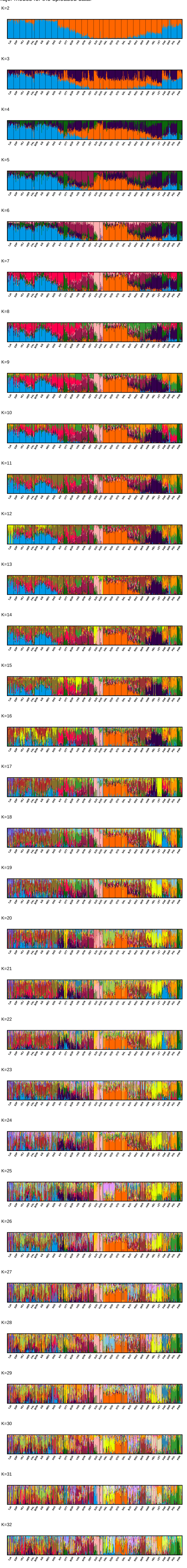


#### Minor modes for the uploaded data:

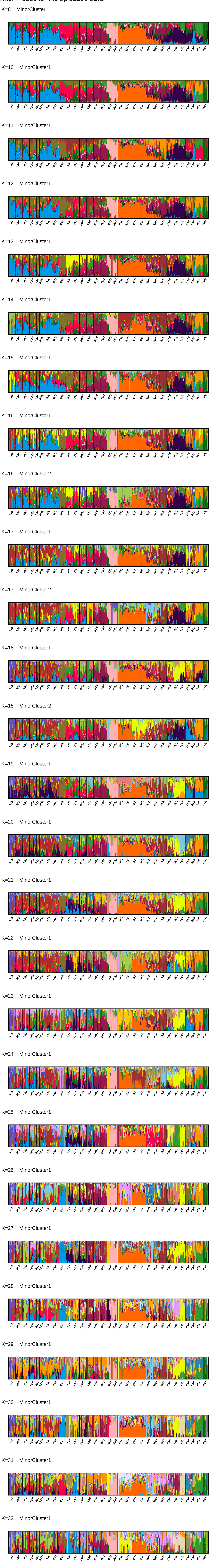


#### Division of runs by mode:

| | |
|---|---|
| K=2 | 10/10 |
| K=3 | 10/10 |
| K=4 | 10/10 |
| K=5 | 10/10 |
| K=6 | 10/10 |
| K=7 | 10/10 |
| K=8 | 8/10, 2/10 |
| K=9 | 10/10 |
| K=10 | 8/10, 2/10 |
| K=11 | 8/10, 2/10 |
| K=12 | 9/10, 1/10 |
| K=13 | 9/10, 1/10 |
| K=14 | 9/10, 1/10 |
| K=15 | 8/10, 2/10 |
| K=16 | 7/10, 2/10, 1/10 |
| K=17 | 7/10, 2/10, 1/10 |
| K=18 | 7/10, 2/10, 1/10 |
| K=19 | 7/10, 3/10 |
| K=20 | 8/10, 2/10 |
| K=21 | 8/10, 2/10 |
| K=22 | 9/10, 1/10 |
| K=23 | 9/10, 1/10 |
| K=24 | 9/10, 1/10 |
| K=25 | 9/10, 1/10 |
| K=26 | 9/10, 1/10 |
| K=27 | 9/10, 1/10 |
| K=28 | 9/10, 1/10 |
| K=29 | 9/10, 1/10 |
| K=30 | 9/10, 1/10 |
| K=31 | 8/10, 2/10 |
| K=32 | 9/10, 1/10 |
